# LOCALE: Local-Alignment Embeddings for Noise-Robust DNA Search at Scale^†∗^

**DOI:** 10.64898/2026.05.12.724581

**Authors:** Ryan P. Synk, Prashant Pandey, S. Cenk Sahinalp, Ramani Duraiswami

## Abstract

Searching petabase-scale repositories of raw sequencing data such as the NIH Sequence Read Archive (SRA) could unlock biological insights that remain inaccessible. Existing sequence-search methods either do not scale well or rely on exact *k*-mer matching that is brittle to genomic variation, which is high both within and across species and is further compounded by sequencing error. We recast sequence search as dense retrieval: we learn sequence embeddings whose inner-product similarity ranks locally aligned sequences above unaligned ones. We introduce LOCALE, an encoder trained with an InfoNCE objective on biologically informed augmentations: overlapping crops of parent sequences corrupted with substitutions, insertions, and deletions. This training recipe, rather than a specialized architecture, consistently induces variation-robust representations across four encoders spanning three tokenization schemes, improving Recall@*R_q_* by 7–39 points at 10% genomic variation. On a 50-accession SRA benchmark LOCALE retains 63.2% Recall@*R_q_* at 10% variation while existing *k*-mer indexes and DNA embedding methods fall below 35%, and on the full 4,571-accession MetaGraph benchmark (296 Gbp, 2.06 billion vectors) it achieves 4.5× the AUPRC of MetaGraph, the state-of-the-art *k*-mer index, with a single-node GPU IVF-PQ index retaining a 3.6× advantage at MetaGraph’s query latency. Our encoder and training strategy generalize to real biological variance: on a cross-genotype Hepatitis B task, querying reads from one genotype against accessions of another with *>*8% natural divergence, LOCALE reaches 0.813 Recall@*R_q_* versus 0.518 for MetaGraph.

## 1 Introduction

Sequence search over a read repository is a dense-retrieval problem with an unusual notion of relevance. A query is a raw sequencing read of 100–300 bp; a document is an assembled sequence of kilobases; the query is relevant to the document if some substring of the document aligns to it at high identity. Relevance is therefore *partial*, since only a substring need match; *asymmetric*, since the query is far shorter than its target; and *variation-invariant*, since the query differs from its match by substitutions, insertions, and deletions. No existing embedding objective targets this relation: DNA sequence embeddings regress *global edit distance* [7, 6, 4, 40, 25], under which a 100 bp exact match inside a 1000 bp sequence has an edit distance of 900 and ranks below unrelated sequences of the query’s own length.

The repository that motivates the problem is the Sequence Read Archive (SRA), whose growth tracks the falling cost of sequencing [38]: it held 47 petabytes in 2025 and grows by 16% a year [34]. Its unit is the *accession*, the reads of one sequencing run, typically one biological sample and often tens of millions of reads. Following repository-scale practice [21, 5], we index assembled contigs, which remove the redundancy of raw reads while preserving each sample’s sequence content, and query with raw reads. The methods that scale to this repository, MetaGraph [21] above all, test exact *k*-mer membership [33, 35], so a single substitution invalidates up to *k* overlapping *k*-mers and indels disrupt more; seed-and-extend aligners [2, 36] tolerate variation but do not scale. Variation is the common case rather than the exception: Hepatitis B virus genotypes differ by more than 8% over the whole genome [31, 30], homologous regions of related species diverge further [17], and legacy long-read data in the SRA carry sequencing error above 10% [9]. A representation for repository search must therefore preserve relatedness after exact identity is lost.

Our key observation is that retrieval does not require an embedding that predicts alignment scores or edit distance; it requires only the correct *ranking*, with locally aligned sequences above unaligned ones. We learn this ranking by contrastive training: a DNABERT-2-style encoder [41] is trained with InfoNCE on positive pairs formed from overlapping crops of one parent sequence, one of which is corrupted with biologically informed substitutions, insertions, and deletions (Figure 1). The inner product of the resulting embeddings ranks aligned pairs above unaligned ones, which turns sequence search into dense retrieval over billions of vectors, with the approximate-nearest-neighbor tooling that comes with it.

**Figure 1:**
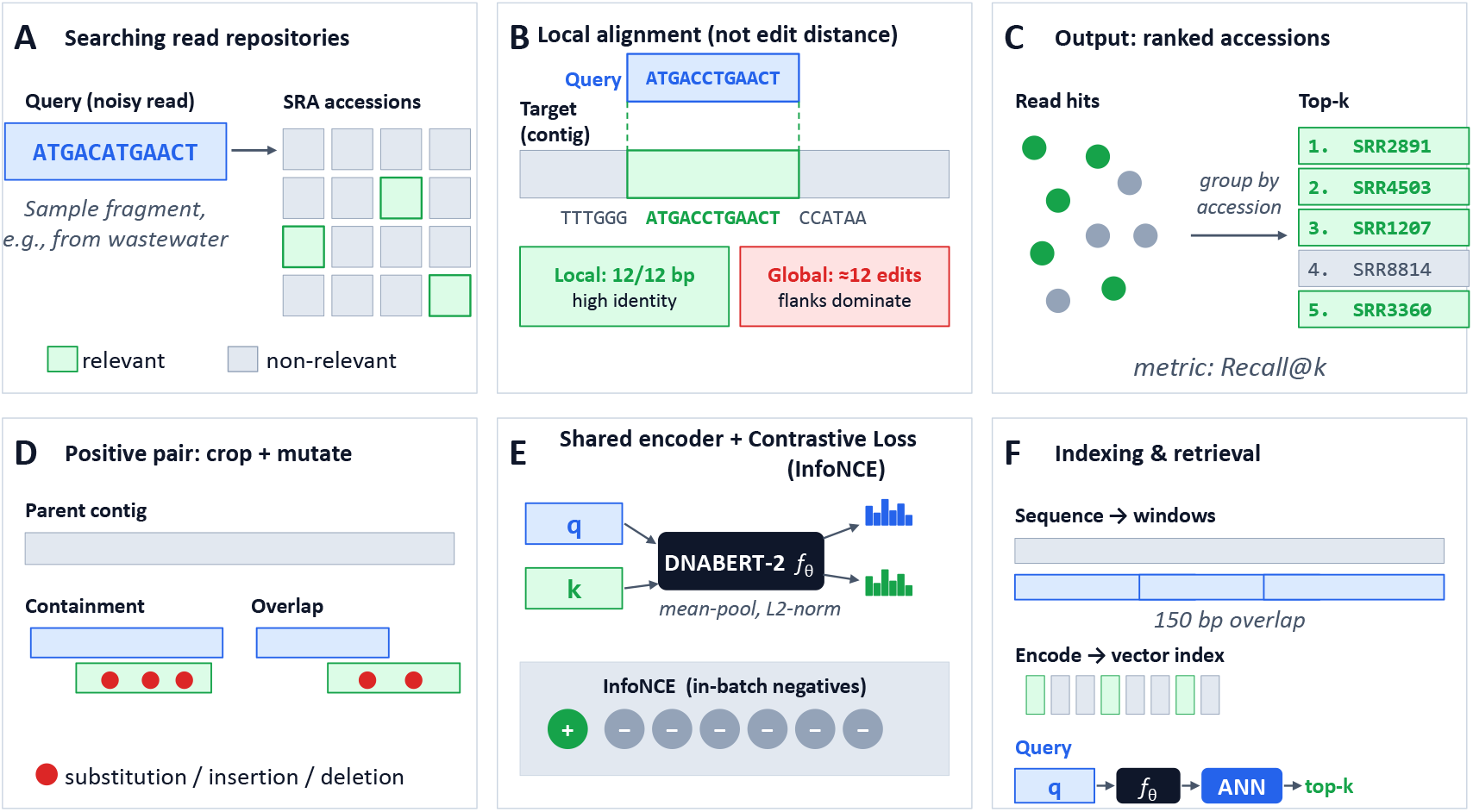
Overview of LOCALE. (Top row: problem.) **(A)** A read carrying genomic variation is queried against SRA accessions, only some of which are relevant. **(B)** Relevance is *local alignment* : a high-identity match to a substring of a target (green) counts even when global edit distance is dominated by unmatched flanks (red). **(C)** Sequence-level hits are grouped by accession, each accession is scored by its best hit, and accessions are returned ranked. **(Bottom row: method.) (D)** Positive pairs are containment or overlap crops of one parent contig, with one crop corrupted by substitutions, insertions, and deletions. **(E)** Both crops pass through a shared encoder *f_θ_* trained with InfoNCE (§3). **(F)** At index time, contigs are split into overlapping windows and embedded; at query time, the query is embedded and its nearest windows are retrieved by inner product (§3).

Our central empirical finding is that *robustness is a property of the training recipe, not of the encoder*. Without variation-augmented contrastive fine-tuning, encoders retain as little as 15% Recall@*R_q_* at 10% variation (Table 3); the same objective and augmentations, applied to four encoders spanning byte-pair, non-overlapping *k*-mer, and single-nucleotide tokenization, raise Recall@*R_q_* by 7–39 points in every case, and a 50M-parameter encoder then matches a 117M-parameter one. The robustness transfers beyond the synthetic corruption used in training: querying reads of one Hepatitis B genotype against accessions of another, LOCALE reaches 0.813 Recall@*R_q_* against 0.518 for MetaGraph and 0.129 for a prior DNA embedding model.

**Table 1:** 50-accession benchmark: Latency is wall-clock for 500-query batch (mean of 4 timed repeats after warm-up). MetaGraph and MMseqs2 run on CPU, exact dense search on 4×A100, and IVF-PQ (1,360 lists, 128×8-bit PQ with fp32 rerank of the top 10 hits per chunk) on a single A100. Full values with standard errors in Table 11.

| Method | AUPRC |  |  | Latency<br>(s) | Index<br>(GB) |
| --- | --- | --- | --- | --- | --- |
|  | 0% | 5% | 10% |  |  |
| **LOCALE** | 0.949 | **0.889** | **0.708** | 2.26 | 30.8 |
| **LOCALE + IVF-PQ** | 0.861 | 0.777 | 0.592 | 0.76 | 1.38 |
| ESA | 0.953 | 0.688 | 0.360 | 2.66 | 40.8 |
| LLM-ED | 0.836 | 0.640 | 0.448 | 2.24 | 30.8 |
| MetaGraph | **0.983** | 0.727 | 0.159 | **0.22** | **0.38** |
| MMseqs2 (alignment reference) | 1.000 | 0.997 | 0.951 | 10.9 | 18.8 |

**Table 2:** Scaling. MetaGraph vs. LOCALE at 50 accessions (1.6 Gbp, 10.7M vectors), 500 (23 Gbp, 153M) and the full 4,571-accession benchmark (296 Gbp, 2.06B). Index size is on disk; for MetaGraph at 4,571 it is the sum of 8 shards. IVF-PQ is the cuVS GPU index (128×8-bit product quantization, 136 B per vector, fp32 rerank of the top 10 hits per chunk) with *m* = 100. Standard errors ≤ 0.02; full values with error bars in Tables 11–13.

| Accessions | Method | Index<br>(GB) | Avg Recall@$R_q$ | | | AUPRC | | |
| --- | --- | --- | --- | --- | --- | --- | --- | --- |
|  |  |  | 0% | 5% | 10% | 0% | 5% | 10% |
| 50 | MetaGraph | 0.38 | **0.982** | 0.714 | 0.113 | **0.983** | 0.727 | 0.159 |
|  | LOCALE (fp32) | 30.8 | 0.925 | **0.850** | **0.632** | 0.949 | **0.889** | **0.708** |
|  | LOCALE (IVF-PQ) | 1.38 | 0.823 | 0.727 | 0.517 | 0.861 | 0.777 | 0.592 |
| 500 | MetaGraph | 4.04 | **0.954** | 0.654 | 0.097 | **0.963** | 0.659 | 0.106 |
|  | LOCALE (fp32) | 438 | 0.876 | **0.764** | **0.458** | 0.901 | **0.796** | **0.507** |
|  | LOCALE (IVF-PQ) | 19.6 | 0.810 | 0.675 | 0.400 | 0.835 | 0.704 | 0.437 |
| 4,571 | MetaGraph | 36.2 | **0.892** | 0.556 | 0.076 | **0.920** | 0.569 | 0.080 |
|  | LOCALE (fp32) | 5,894 | 0.803 | **0.671** | **0.323** | 0.847 | **0.707** | **0.357** |
|  | LOCALE (1-bit RaBitQ) | 200 | 0.794 | 0.638 | 0.265 | 0.837 | 0.678 | 0.292 |
|  | LOCALE (IVF-PQ) | 263 | 0.748 | 0.600 | 0.275 | 0.781 | 0.620 | 0.287 |

**Table 3:** Backbone generalization. Average Recall@*R_q_* on the 50-accession benchmark at three evaluation variation rates, ± bootstrap standard error over 500 queries. Every row is fine-tuned on the same 6M Logan contig pairs with the same objective and augmentation ladder; “heavy” is the augmentation configuration of LOCALE. DNABERT-2 uses the 5,859-step schedule of §3; the other backbones use 11,718 steps at matched peak learning rate.

| Backbone | Aug. | 0% | 5% | 10% |
| --- | --- | --- | --- | --- |
| DNABERT-2 | none | 0.948±0.007 | 0.755±0.016 | 0.273±0.017 |
| | heavy | 0.927±0.009 | 0.862±0.012 | **0.662±0.018** |
| NT-50M | none | 0.952±0.007 | 0.854±0.013 | 0.539±0.019 |
| | heavy | 0.935±0.008 | 0.865±0.012 | **0.684±0.018** |
| HyenaDNA | none | 0.940±0.008 | 0.843±0.013 | 0.538±0.019 |
| | heavy | 0.911±0.010 | 0.818±0.014 | **0.605±0.018** |
| ESA (retrained) | none | 0.920±0.010 | 0.512±0.019 | 0.153±0.013 |
| | heavy | 0.898±0.010 | 0.638±0.018 | **0.302±0.017** |

**Table 4:**
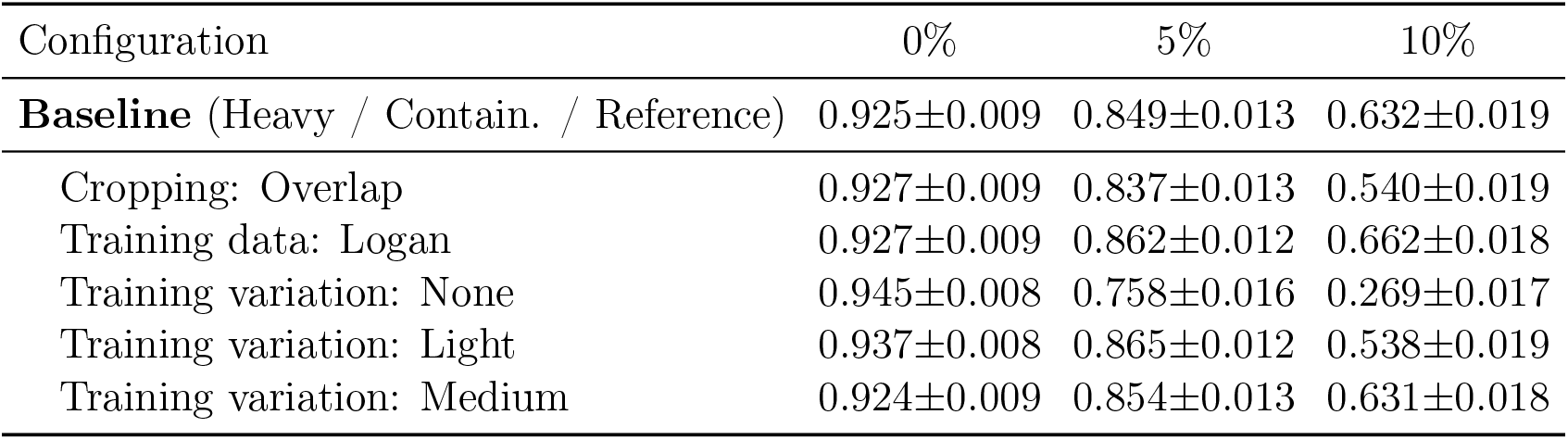
Ablations. Average Recall@*R_q_* on the 50-accession benchmark at three evaluation variation rates, ± bootstrap standard error. **Baseline**: heavy training variation, containment cropping, reference-genome training data (the released LOCALE model). Each row below the baseline changes one axis.

Our main contributions are:

- **Formulate sequence search as variation-robust dense retrieval,** with partial, asymmetric relevance defined by local alignment, and learn it as a ranking rather than a regression.
- Show that robustness comes from the objective and augmentation, not the backbone: our recipe raises 10%-variation Recall@*R_q_* by 7–39 points across four encoders and three tokenizers.
- **Demonstrate transfer from synthetic to natural variation** on cross-genotype Hepatitis B reads with more than 8% divergence.
- **Scale to the full 4,571-accession MetaGraph benchmark** (2.06 billion vectors), where a single-node GPU index matches the *k*-mer index’s query latency at 3.6× its AUPRC, and release the benchmark.

## 2 Related Work

### Seed-and-extend

BLAST [2] and MMseqs2 [36] seed on exact *k*-mers and extend by dynamic programming. Their seed index is a flat map from *k*-mers to positions, so query time grows linearly with the database, and they target reference genomes rather than read repositories. Since our relevance labels are themselves alignments, we use MMseqs2 as the alignment-quality reference, not as a scalable competitor.

### Experiment discovery

To scale to billions of reads, *k*-mer indexes recast search as *experiment discovery*: return the accessions containing at least a set fraction of the query’s *k*-mers. Sequence Bloom trees [35] and Mantis [33] made this practical over thousands of experiments, colored de Bruijn graphs [1] compress the *k*-mer-to-accession map, and MetaGraph [21] scales it to a large fraction of the SRA; Logan-Search [5] and Fulgor [10] rest on the same exact-*k*-mer foundation. All inherit the brittleness of exact *k*-mer membership under variation (§1). Additionally, figure 2 shows that *k*-mers are susceptible to false positives, since high *k*-mer overlap does not imply alignment. We take MetaGraph as their representative: it is the most scalable published method, it indexes the same Logan contigs we do, and we evaluate on its own benchmark accessions (§4).

**Figure 2:**
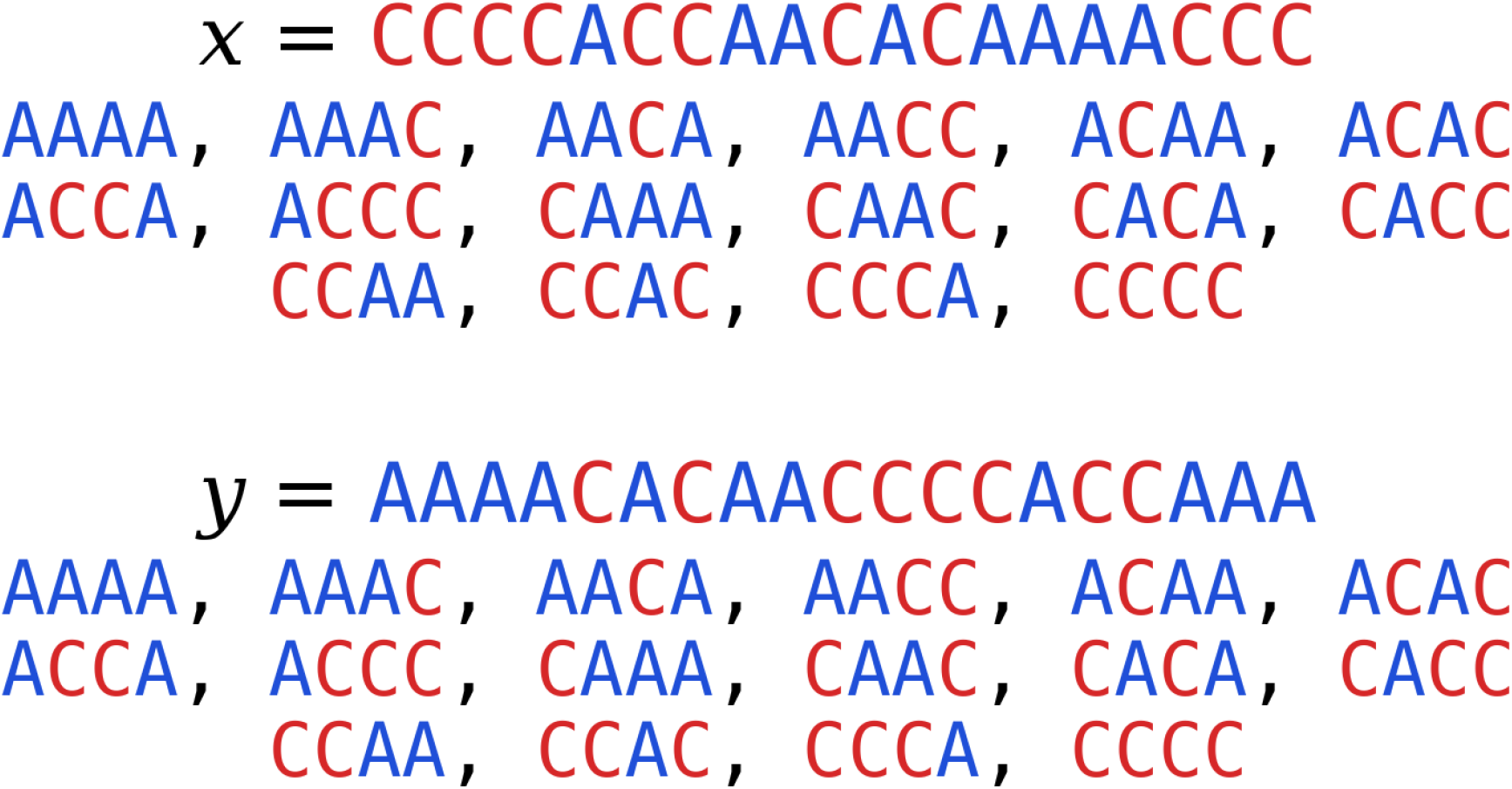
Sequences *x* and *y* have the same *k*-mer representation but low sequence similarity. *k*-mer-based representations employed in MetaGraph and Mantis are not the right proxy for sequence search since they can introduce a large number of false positives.

### DNA sequence embeddings

Metric-learning methods embed sequences so that vector distance approximates global edit distance [7, 6, 4, 40, 25], and some require fixed-length inputs [7, 40]. Because edit distance is dominated by the length difference between a read and a contig window, such models rank globally similar but locally irrelevant sequences above true local matches (§4). Embed-Search-Align [15] and NeurALigner [3] embed reads to align them to a single reference genome and train mainly on high-identity pairs; we address the inverse problem, indexing many accessions and returning the matching ones, and adapt Embed-Search-Align as a baseline, whose retrieval degrades as variation grows (§4).

### Dense retrieval

Dense retrieval [22, 16, 12] maps queries and documents into a shared space, often with contrastive objectives over augmented views [16, 14], and pairs the encoder with approximate nearest-neighbor indexes [19] to search billions of documents under lexical variation. Scale, tolerance to variation, and sublinear query time are exactly what experiment discovery over raw reads requires; LOCALE is a dense retriever whose relevance is local alignment.

## 3 Method

We learn an encoder *f_θ_* : Σ^∗^ → ℝ*^D^* over Σ = {*A, C, G, T* } such that the inner product *f_θ_*(*s*)^⊤^*f_θ_*(*p*) ranks locally aligned sequences *s, p* above unaligned ones. This section gives the objective, the construction of positive pairs, the encoder and training data, and the search procedure.

### Training objective

Given a batch of *B* positive pairs of aligned sequences 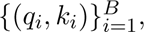 we minimize the InfoNCE loss [37] with in-batch negatives,

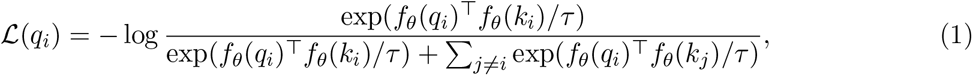

with temperature *τ* = 0.05 throughout. The loss pushes each positive pair together and every other pair in the batch apart; it constrains the ranking of similarities, not their values.

### Data augmentation

Positive pairs are two crops of one parent sequence, one of which is then corrupted (Figure 1). A *containment* crop pairs a sequence with a subsequence contained inside it; an *overlap* crop pairs two sequences that overlap at one end without either containing the other. Crop lengths are uniform between 31 and 256 bp, and the shorter crop must cover at least 40% of the longer one (for overlap crops, the overlapping region must span at least 40% of the longer crop). Containment is the relation between a read and the contig it aligns to, and the ablations in §4.2 favor it. We then inject variation into one crop: a target identity *ρ* ∈ [0*, ρ*_max_] is drawn from a Beta distribution parametrized, following Wick [39], by its mean and standard deviation rather than by *α, β*; a crop of length *L* receives *n* = ⌊*L*(1 − *ρ*)⌋ mutations at positions sampled without replacement within the aligned region, each independently a substitution, insertion, or deletion with probabilities 0.4, 0.3, 0.3. Substitutions draw uniformly from the three other bases and insertions place a uniform random base before the sampled position. Which crop is mutated, and which serves as anchor *q_i_* versus key *k_i_* in Eq. 1, are both chosen uniformly at random. We use four augmentation strengths, *none*, *light*, *medium*, and *heavy* ; the final model uses heavy, with identity mean 80%, *ρ*_max_ = 88%, and standard deviation 6 points, so a typical training pair differs at about one base in five. Light and medium settings are in Table 7.

### Encoder

We use DNABERT-2 [41] (117M parameters, byte-pair tokenizer with a 4096-token vocabulary, pretrained on 700 bp sequences) and fine-tune all parameters under Eq. 1. Token representations *h_t_* ∈ ℝ*^D^*, *D* = 768, are mean-pooled over non-padding positions and *L*_2_-normalized, with cosine similarity on the unit sphere. Optimizer and schedule are in Appendix A.3.1.

### Training data

We train on two corpora with the same pipeline and budget, 6 million pairs each. The *Logan* corpus is contigs from the Logan project [5], which assembled every accession in a Dec. 31, 2025 snapshot of the SRA; we subsample the 5,056-accession MetaGraph benchmark slice [21] to 50 training accessions, disjoint from every accession used in evaluation (species breakdown in Table 8). The *reference* corpus is the NCBI RefSeq [13] genomes of the species in those 50 accessions, chunked into non-overlapping 1024 bp parents (Appendix A.2.4). The released model uses the reference corpus; §4.2 shows the two perform similarly.

### Indexing and search

For accessions U = {*A*_1_*, . . ., A_N_* }, each a set of sequences, we embed every sequence into a vector index, splitting sequences longer than the encoder context into 256 bp windows with 150 bp overlap. A query *q* and its reverse complement are both embedded, the top-*m* vectors by inner product are retrieved for each, and the union N*_m_*(*q*) is aggregated to accessions by the best hit,

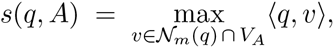

where *V_A_* are the vectors of accession *A*; accessions are returned in descending *s*(*q, A*). We use *m* = 100 at 50 and 500 accessions and *m* = 1000 for exact search at 4,571, where the two agree within 0.5 points of Recall@*R_q_* (0.798 vs. 0.803 without variation; full sweep in Appendix B). The index is an ordinary vector index, so quantization such as RaBitQ [11] and inverted-file search apply unchanged; §4.4 uses both.

## 4 Experiments

### Problem Definition and Metrics

We evaluate LOCALE’s ability to retrieve accessions containing local matches to query sequences with genomic variation using a benchmark constructed from real SRA data. Given an accession universe U and query *q*, an accession *A* is relevant if *q* has a local alignment to a sequence in *A* with identity at least *ι*, where identity is the fraction of exact base-pair matches in the aligned region. We set *ι* = 0.9. A method returns accessions in ranked order; for the top-*k* retrieval set ℝ*_k_*(*q*) and relevant set A*_q_*, we define Recall@*k*(*q*) = |A*_q_* ∩ ℝ*_k_*(*q*)|*/*|A*_q_*|. We report Average Recall@*R_q_*, where *R_q_* = |A*_q_*|, averaging Recall@*R_q_* over queries; this evaluates recall at the exact number of relevant accessions and equals 1 for perfect retrieval. We also report AUPRC, which summarizes the precision-recall tradeoff over all *k*.

### Dataset

We evaluate on the same accessions as the MetaGraph benchmark [21], which we refer to as the 100-studies dataset. It contains 5,056 accessions spanning 100 SRA studies. For each accession, we index contigs assembled from its raw reads by Logan [5]. We evaluate at three scales: 50 accessions (1.6 Gbp, 10.7M vectors), 500 accessions (23 Gbp, 153M vectors), and 4,571 accessions (296 Gbp, 2.06B vectors). The full test set excludes contigs used for training and contigs not yet available in Logan; the two smaller sets are sampled proportionally to the organism distribution of the full dataset (Appendix §A.2.2). No accession overlaps between model training and evaluation.

### Query Sourcing

Following MetaGraph [21], we draw queries from raw reads of the indexed accessions. This provides natural ground truth: a read is, with high probability, locally alignable to a contig from its source accession. Retrieval remains non-trivial because Logan contigs are assembled from many reads and each read represents only a local match.

### Relevance labeling and filtering

We establish ground-truth relevance by aligning each query against every indexed accession with Minimap2 [24], labeling accession *A* relevant if any alignment has identity ≥ *ι* = 0.9. Let A*_q_* denote the resulting relevant set. We discard queries whose source accession is not in A*_q_*, since such reads are not alignable to their source contigs (e.g., due to read-quality or contig-cleaning artifacts), and queries for which |A*_q_*| exceeds 20% of the index, as highly conserved or repetitive queries yield high recall even under random ranking. We uniformly sample reads from accessions, apply these filters, and fix 500 queries for the 50- and 500-accession benchmarks and 1,000 for the 4,571-accession benchmark. Queries are stored without strand normalization; 51–55% align to the reverse strand of their source contig (Table 19).

### Simulated genomic variation

To evaluate retrieval under genomic variation, we generate two variants of each query with Mutation-Simulator [23], using nominal SNP rates of 5% and 10% and insertion/deletion rates equal to 10% of the SNP rate. LOCALE is trained with the augmentation procedure described above, not Mutation-Simulator, so these variants represent an unseen corruption process. Variants are generated once and fixed, ensuring byte-identical inputs across methods. We refer to the 5% and 10% settings by their nominal SNP rates, corresponding to target identities of approximately 95% and 90%, respectively. For each variant, we retain the relevant set A*_q_* computed from the original unmutated read, thereby measuring whether a method can recover the original relevant accessions after query corruption.

We evaluate up to 10% variation to test robustness at substantial genomic divergence, spanning the within-species and cross-strain variation motivating our benchmark and reaching our relevance threshold *ι* = 0.9. This also captures regimes where sequencing errors can compound biological variation in legacy SRA data [9].

### Baselines

We compare LOCALE to:

- **MetaGraph** [21], the state-of-the-art *k*-mer index and the one that scales to the largest fraction of the SRA (§2), with accessions scored by *k*-mer overlap percentage. We run it on its own benchmark accessions and on the same Logan contigs it indexes at SRA scale. Following their paper, we build indexes with *k* = 31 and query with a *k*-mer discovery fraction of 0.0.
- **MMseqs2** [36], a seed-and-extend method, with accessions ranked by best alignment bit score.
- Two vector embedding models, retrieved via the same accession-level aggregation as LOCALE: **LLM-ED** [25], an edit-distance model; and **Embed-Search-Align** [15], a read-to-reference genome mapping model.

For embedding baselines, we substitute their encoder into our indexing and retrieval pipeline (§3). MMseqs2 is an alignment-based reference point rather than a scalable competitor: its index grows to 198 GB at 500 accessions (49× MetaGraph’s) and a 500-query batch takes 80 s on a 128-core CPU node, so we run it at 50 and 500 accessions but not on the full 4,571-accession benchmark. Dense search runs on 4×A100 nodes and MetaGraph/MMseqs2 on 128-core CPU nodes; full hardware in Appendix D.1.

### 4.1 Retrieval Quality

Figure 3 reports Average Recall@*R_q_* versus variation rate on the 50-accession dataset; Recall@*k* at 10% variation is shown in Figure 5. At 10% variation, LOCALE retains 63.2% Average Recall@*R_q_*, compared with 25.1% for ESA, 34.3% for LLM-ED, and 11.3% for MetaGraph. Without variation, all exhaustive-search methods except LLM-ED exceed 92%, showing that the benchmark becomes discriminative only under variation. ESA matches LOCALE without variation (93.6% vs. 92.5%) but degrades sharply with increasing variation, reflecting its lack of variation-augmented training. LLM-ED trails LOCALE even without variation (78.5% vs. 92.5%), consistent with regressing exact edit distance rather than learning the ranking needed for retrieval. MetaGraph is particularly brittle: 25.0% of queries return no results at 5% variation and 88.4% at 10%, as too few 31-mers survive; an accession never retrieved receives no credit at any *k*. Table 1 shows the same ordering under AUPRC. Without variation, LOCALE, ESA, and MetaGraph are within 0.04 AUPRC (0.949, 0.953, and 0.983). At 10% variation only LOCALE remains robust: it retains 0.708 AUPRC, compared with 0.360 for ESA, 0.448 for LLM-ED, and 0.159 for MetaGraph. MMseqs2 remains above 0.95 at all variation rates, indicating substantial headroom between learned retrieval and exhaustive alignment (0.708 vs. 0.951).

**Figure 3:**
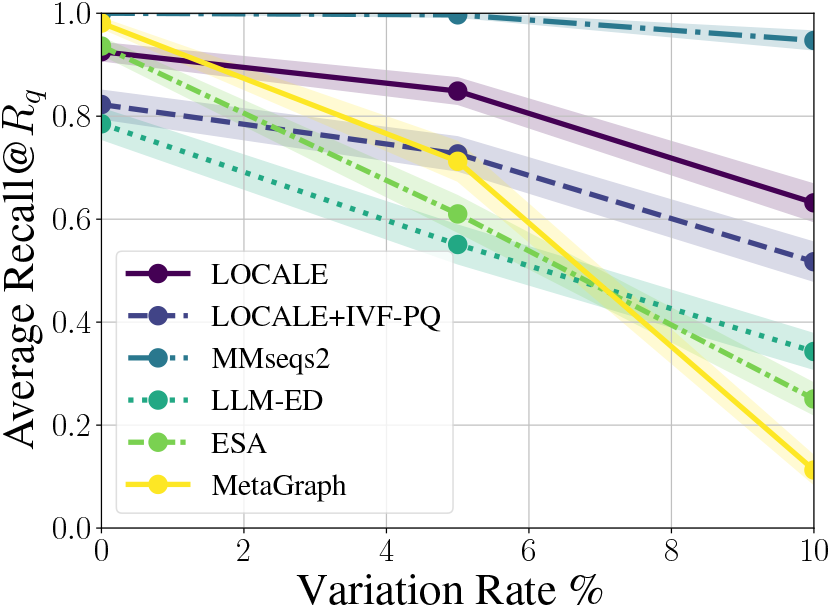
Average Recall@*R_q_* over the 50-accession dataset, averaged across 500 queries. Nominal SNP rates of 0%, 5% and 10% correspond to query identities of 100%, 95% and 90%.

**Figure 4:**
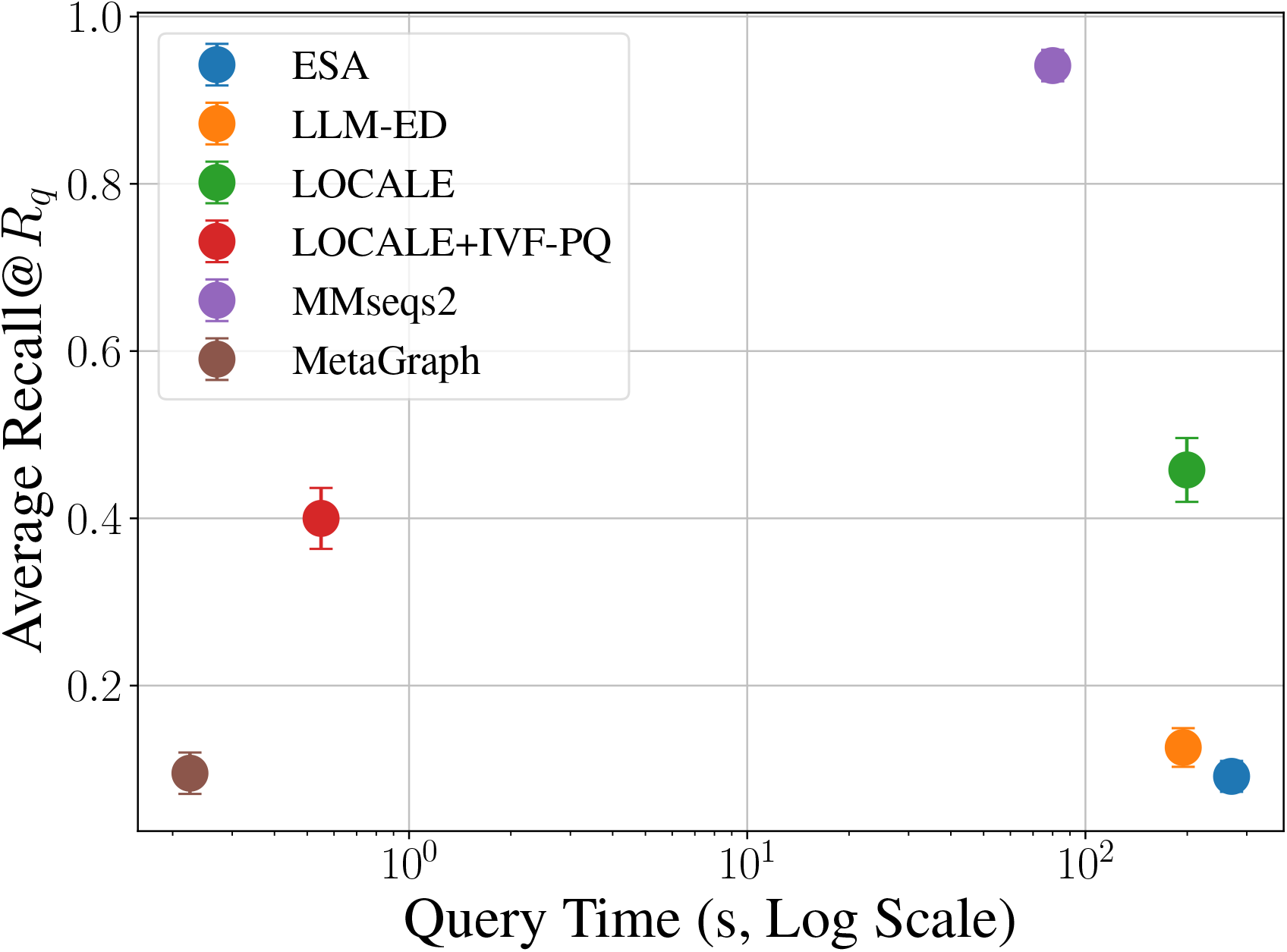
Recall@*R_q_* vs. query time (log scale) for a 500-query batch on the 500-accession benchmark at 10% variation; upper-left is better. Dense methods run on one 4×A100 node, MetaGraph and MMseqs2 on one CPU node. Exact float32 search streams the 438 GB index from disk on every pass, so its time is I/O-bound; the IVF-PQ index (19.6 GB, GPU-resident) answers the same batch in 0.55 s at a cost of 6 points of recall, and still retrieves 4× more than MetaGraph.

**Figure 5:**
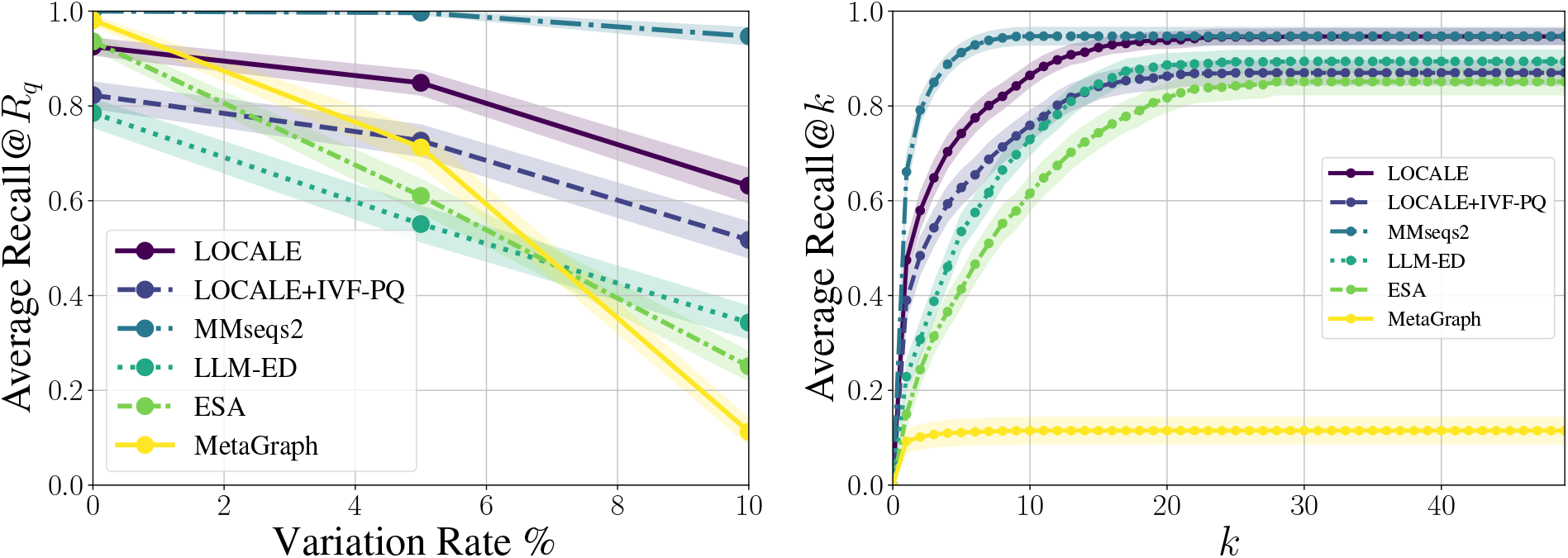
Left: Average Recall@*R_q_* over the 50-accession dataset, averaged across 500 queries. Nominal SNP rates of 0%, 5% and 10% correspond to query identities of 100%, 95% and 90%. Right: Average Recall@*k* for increasing numbers of returned accessions *k* (out of 50) under 10% variation. MetaGraph’s curve saturates at *k* ≈ 2 because an accession it never returns earns no credit at any *k*.

The timing and index sizes reveal the corresponding efficiency tradeoff. On 50 accessions, MMseqs2 takes 10.9 s for 500 queries, versus 2.26 s for exact LOCALE and 0.76 s for LOCALE IVF-PQ, while its index reaches 198 GB at 500 accessions. MetaGraph is fastest (0.22 s) and smallest (0.38 GB), but its AUPRC at 10% variation is only 0.159. LOCALE IVF-PQ is 3.5× slower and 3.6× larger than MetaGraph, but achieves 3.7× higher AUPRC (0.592 vs. 0.159); §4.4 shows that inverted-file indexing further narrows the latency gap at full scale.

### 4.2 What Drives Robustness to Genomic Variation?

#### Encoder Backbones

Table 4 shows that increasing augmentation improves recall on queries with genomic variation. We next ask whether this robustness comes from the training recipe or from the choice of encoder and tokenizer. We apply the same fine-tuning procedure to Nucleotide Transformer (NT-50M, non-overlapping *k*-mers [8]), HyenaDNA (single-nucleotide tokens [29]), and Embed-Search-Align (ESA). Each model is trained on the same 6M Logan contig pairs, with identical augmentation and training budgets, with and without variation augmentation. The DNABERT-2 rows in Table 3 are trained in the same sweep, so all models share the same corpus and recipe.

#### The training recipe generalizes across backbones

Heavy augmentation consistently improves Recall@*R_q_* at 10% variation: by 39 points for DNABERT-2, 15 for Nucleotide Transformer, 7 for HyenaDNA, and 15 for ESA. Increasing augmentation also reduces clean-query recall by at most 3 points. Thus, variation robustness is consistently induced by the training recipe rather than tied to a particular backbone or tokenizer.

#### Robustness does not require a large encoder

Nucleotide Transformer (50M parameters) matches DNABERT-2 (117M) at 10% variation (0.684 vs. 0.662, within one standard error). This further indicates that the training recipe, rather than encoder capacity, is the primary driver of robustness.

#### Ablation Studies

Table 4 isolates the effects of individual training choices. Cropping strategy matters primarily under variation: containment cropping improves Recall@*R_q_* by 9.2 points at 10% variation (0.632 vs. 0.540), while the two are indistinguishable without variation. Containment pairs reflect the deployment relationship between a read and its matching contig, since one crop is a substring of the other; because anchor and key roles are randomized (§3), the improvement is not due to directional asymmetry in the loss.

Training on Logan contigs performs comparably to curated reference genomes (0.662 vs. 0.632 at 10% variation; 0.862 vs. 0.849 at 5%, both within roughly one standard error), showing that assembled metagenomic contigs provide sufficient diversity for contrastive pretraining despite being less curated. This is practically useful because Logan provides orders of magnitude more sequence than curated references.

Finally, augmentation strength directly controls robustness with little cost on clean queries. Recall@*R_q_*at 10% variation increases from 0.269 with no variation augmentation to 0.538, 0.631, and 0.632 for light, medium, and heavy augmentation, respectively, while clean-query recall remains within a 2-point range (0.924–0.945). Thus, stronger augmentation yields a 36-point robustness gain with at most a 2-point clean-data loss.

### 4.3 Retrieval Across Real Biological Divergence

#### Cross-genotype generalization

To test whether robustness to synthetic variation transfers to genuine biological divergence, we evaluate cross-genotype retrieval on Hepatitis B Virus (HBV), whose genotypes differ by at least ∼8% across the whole genome [31, 30]. We select SRA runs by metadata and verify them against reference genomes (Appendix A.3.2); query and target runs come from different BioProjects to avoid BioProject-specific matches. We use reads from 10 genotype-D runs as queries and index 5 genotype-B accessions plus the 50-accession benchmark as distractors, for 55 accessions total. The five genotype-B accessions are relevant to every query (*R_q_* = 5), while genotype-D source accessions are excluded, ensuring that all credited retrievals are cross-genotype. We compare methods deployable at repository scale and include MMseqs2 as an alignment-quality reference.

As a sanity check on the degree of divergence, among 5,000 reads from genotype-D run SRR12535932, 84.4% have a 31-mer match to a genotype-D reference, versus 49.2% to a genotype-B reference. Table 5 shows that LOCALE achieves 0.813 Recall@*R_q_*, compared with 0.518 for Meta-Graph, 0.316 for LLM-ED, and 0.129 for ESA; its single-GPU IVF-PQ index also exceeds MetaGraph (0.714 vs. 0.518). MetaGraph returns no result for 22.8% of queries because no 31-mers survive. These results show that robustness learned from synthetic corruption transfers to naturally occurring cross-genotype divergence.

**Table 5:** Cross-genotype HBV retrieval. Genotype-D reads queried against indexed genotype-B accessions (*R_q_* = 5) among 50 distractor accessions, relevance from curated genotype labels; ± is the bootstrap standard error over 500 queries. “No result” is the fraction of queries for which the method returns an empty ranking. MMseqs2 is included as the alignment quality ceiling, consistent with its role in Table 1.

| Method | Recall@$R_q$ | Recall@7 | AUPRC | No result |
| --- | --- | --- | --- | --- |
| **LOCALE** | **0.813**±0.013 | **0.847**±0.012 | **0.850**±0.011 | 0.0% |
| LOCALE + IVF-PQ | 0.714±0.015 | 0.743±0.015 | 0.757±0.014 | 0.0% |
| MetaGraph | 0.518±0.019 | 0.518±0.019 | 0.562±0.017 | 22.8% |
| LLM-ED | 0.316±0.014 | 0.361±0.015 | 0.387±0.014 | 0.0% |
| Embed-Search-Align | 0.129±0.011 | 0.133±0.011 | 0.206±0.010 | 0.0% |
| MMseqs2 (reference) | 0.915±0.010 | 0.915±0.010 | 0.923±0.009 | 3.0% |

### 4.4 Scalability

We evaluate LOCALE at 500 accessions (153M vectors) and the full 4,571-accession benchmark (2.06B vectors). A float32 index for the full benchmark requires 5.9 TB, so we evaluate two compressed indexes: 1-bit RaBitQ [11], searched exhaustively, and GPU IVF-PQ using cuVS [20, 32]. RaBitQ reduces the full index to 200 GB, while IVF-PQ occupies 263 GB on disk and probes 1.7% of lists per query (configuration in Table 13). Tables 2 and 6 compare these indexes with MetaGraph across scales and at full scale, respectively. We omit other embedding baselines at 4,571 accessions because LOCALE outperforms them at 50 and 500 accessions (Tables 1 and 12). MMseqs2 is likewise limited to 500 accessions; extrapolating its measured 8.7 GB-per-Gbp index size to the full benchmark yields roughly 2.5 TB.

Compression substantially reduces storage with modest quality loss. At full scale, 1-bit RaBitQ shrinks the float32 index by 29× (5.9 TB to 200 GB), reducing Recall@*R_q_*by 0.9 points without variation and 5.8 points at 10% variation (0.323 vs. 0.265). IVF-PQ loses 5.5 and 4.8 points, respectively, while float32 reranking recovers 5 points of AUPRC at 10% variation (0.287 vs. 0.236). Both remain substantially more robust than MetaGraph: RaBitQ achieves 3.5× its Recall@*R_q_* and 3.7× its AUPRC at 10% variation (0.292 vs. 0.080), while IVF-PQ achieves 3.6× both metrics. At 500 accessions, IVF-PQ similarly reaches 0.437 AUPRC versus 0.106 for MetaGraph at 10% variation, using 19.6 GB versus 4.0 GB.

IVF-PQ also makes search practical at full scale (Table 6). Exhaustive search over 2.06B vectors takes minutes per 1,000-query batch even with 1-bit vectors and four GPU nodes, whereas IVF-PQ answers the batch in 3.2 s on one GPU node, including embedding and reranking. MetaGraph takes 3.8 s for the same batch on one CPU node as a single request, or 0.46 s with 64 concurrent requests. Thus, LOCALE approaches the latency of the scalable *k*-mer index while retaining 3.6× its AUPRC under 10% variation, although the comparison uses different hardware. IVF-PQ additionally requires 1–3 minutes to load 304 GB of PQ codes into GPU memory before querying. More scalable graph indexes [27, 18, 28] and coarser probing are natural directions for larger accession counts.

**Table 6:** Search cost on the full 4,571-accession benchmark for a 1,000-query batch, both strands embedded. Times are wall-clock seconds at 0% / 5% / 10% variation, mean of 4 timed repeats after warm-up, except the two exact-search rows marked †, which are Slurm job elapsed times for a 4-node run including query embedding, sharded scan, merge and regroup. GPU nodes have 4×A100 (80 GB for IVF-PQ, 40 GB otherwise); CPU nodes have 128 cores. MetaGraph is queried over its 8 shards either as one request (served by one core) or as 64 concurrent requests. AUPRC at 10% variation from Table 13.

| Method | Index (GB) | Hardware | Batch time (s) | AUPRC 10% |
| --- | --- | --- | --- | --- |
| MetaGraph, 1 request | 36 | 1 CPU node | 3.81 / 3.44 / 3.98 | 0.080 |
| MetaGraph, 64 parallel requests | 36 | 1 CPU node | **0.46 / 0.41 / 0.34** | 0.080 |
| LOCALE fp32, exact | 5,894 | 4 GPU nodes | ~735<sup>†</sup> | **0.357** |
| LOCALE 1-bit RaBitQ, exact | 200 | 4 GPU nodes | ~168<sup>†</sup> | 0.292 |
| LOCALE CPU IVF-RaBitQ (FAISS) | 215 | 1 CPU node | 133 / 145 / 139 | 0.332 |
| LOCALE GPU IVF-PQ, no rerank | 263 | 1 GPU node | 3.10 / 2.86 / 2.89 | 0.236 |
| LOCALE GPU IVF-PQ + fp32 rerank | 263 | 1 GPU node | 3.22 / 3.29 / 3.80 | 0.287 |

## 5 Conclusion

We presented LOCALE, a contrastive embedding model that recasts sequence search over raw-read repositories as dense retrieval rather than edit-distance regression: retrieval requires only that aligned pairs score higher than unaligned ones, not that distances be approximated accurately. A simple crop-and-corrupt training recipe confers robustness to genomic variation across four encoder backbones, transfers from synthetic corruption to real cross-genotype divergence, and scales to 4,571 accessions and 2.06 billion vectors, where a compressed LOCALE index trades memory for robustness: 5.5–7.3× MetaGraph’s index size for 3.6–3.7× its AUPRC on 10%-divergent queries, at comparable single-node latency. We view this as evidence that the bottleneck for petabase-scale sequence search is no longer scalability alone but tolerance to genomic variation at scale, and we expect future gains to come from the approximate nearest-neighbor, quantization, and graph-based search techniques the dense-retrieval framing makes available rather than from larger encoders.

Several limitations remain. The index is larger than a *k*-mer index even after compression, and the IVF-PQ index must be resident in roughly 300 GB of GPU memory to be served at the latencies we report; learned codebooks or distilled encoders with smaller embedding dimension may be needed at full SRA scale. Contigs longer than the encoder context are embedded as independent windows, discarding long-range structure, and all queries are short reads (100–300 bp); kilobase-scale queries would require chunking and aggregating across chunk-level matches, which we have not evaluated. Finally, we evaluate on a single benchmark slice of the SRA and on variation rates up to 10%; generalization across sequencing platforms, taxonomic distributions, and more divergent queries remains to be tested.

## AI use statement

We did not use generative AI tools for any task in the required-disclosure list: the research questions, hypotheses, methodology, experimental design, implementation, data generation and cleaning, and interpretation of results are the authors’ own. We used large language models for tasks in the recommended-disclosure list: editing the manuscript for readability, drafting and revising portions of the text from the authors’ notes and result tables, formatting references, editing code for experiments, and typesetting result tables from our experiment spreadsheets. Every AI-assisted passage was reviewed and revised by the authors, every number in the text was checked against the underlying result files, and every reference was verified against its publisher or arXiv record. We take responsibility for the final content of this work, including text, claims or artifacts produced with the aid of generative AI.

## Ethics statement

LOCALE accelerates search over publicly available sequencing data and could enable applications such as faster pathogen surveillance, viral variant tracking, and the characterization of biological “dark matter” sequences in metagenomic samples. Because the released artifact is an embedding encoder for retrieval over already-public sequences (not a generative model and not a model trained on private or restricted data) we consider the dual-use risk to be no greater than that of existing widely deployed sequence search tools such as BLAST. We will release the model and benchmark with documentation describing intended use and known limitations.

## Appendix A

### A.1 Glossary of Biological Terms

- **Raw read**: A raw read is a sequence fragment produced directly by a sequencing instrument before assembly, and typically before substantial error correction or polishing.
- **Contig**: A contig is a contiguous stretch of DNA sequence without gaps, assembled from overlapping sequencing reads using direct sequence information.
- **Accession**: An accession, or accession number, is a unique identifier assigned to a database record or unit of molecular data, such as a sequence, sample, study, or run. For the purposes of this paper, we use the term "accession" to refer to a single sequencing run.
- **SRA**: The Sequence Read Archive (SRA) is NCBI’s public repository for high-throughput sequencing data, storing raw sequencing reads and related alignment information.
- **Logan**: Logan is a large-scale assembled sequence resource built from SRA data that transforms short reads into longer contigs.
- **Local Alignment**: Local alignment is the alignment of subsequences to identify the highest-scoring region of similarity between two biological sequences rather than forcing an end-to-end match.
- **Edit distance**: Edit distance is the minimum number of insertions, deletions, and substitutions required to transform one sequence or string into another.
- *k***-mer**: A *k*-mer is a subsequence of length *k* extracted from a longer biological sequence such as a read, contig, or genome.

### A.2 Data and Training

#### A.2.1 Augmentation Parameters

Parameters for light, medium, heavy augmentation distributions: To simulate errors in sequences at training time, we sample identities from a Beta distribution parametrized by a mean and standard deviation rather than conventional shape parameters *α, η* (following the methodology of [39]). Table 7 lists the three augmentation configurations we used in our experiments:

**Table 7:** Augmentation configurations used in our experiments. Sequence identities are sampled from a Beta distribution reparametrized by mean, standard deviation, and maximum (following Wick [39]) rather than by the conventional shape parameters *α, β*. Values are reported as percent identity.

| Configuration | Mean (%) | Std. dev. (%) | Max (%) |
| --- | --- | --- | --- |
| Light | 95.0 | 2.5 | 99.0 |
| Medium | 90.0 | 6 | 98.0 |
| Heavy | 80.0 | 6 | 88.0 |

#### A.2.2 Logan Data

**Table 8** describes the species breakdown of the Logan training dataset. Tables 9 and 10 describe the 50 and 500 acc tests sets, respectively. Each table contains the number of accessions from the respective species, most sets have majority mouse and human data, consistent with their status as the most heavily represented species in the SRA [5]. The tables list the total number of base pairs and total number of contigs across all accessions from the species. They also list n50, which is a metric for identifying sequence length.

**Table 8:**
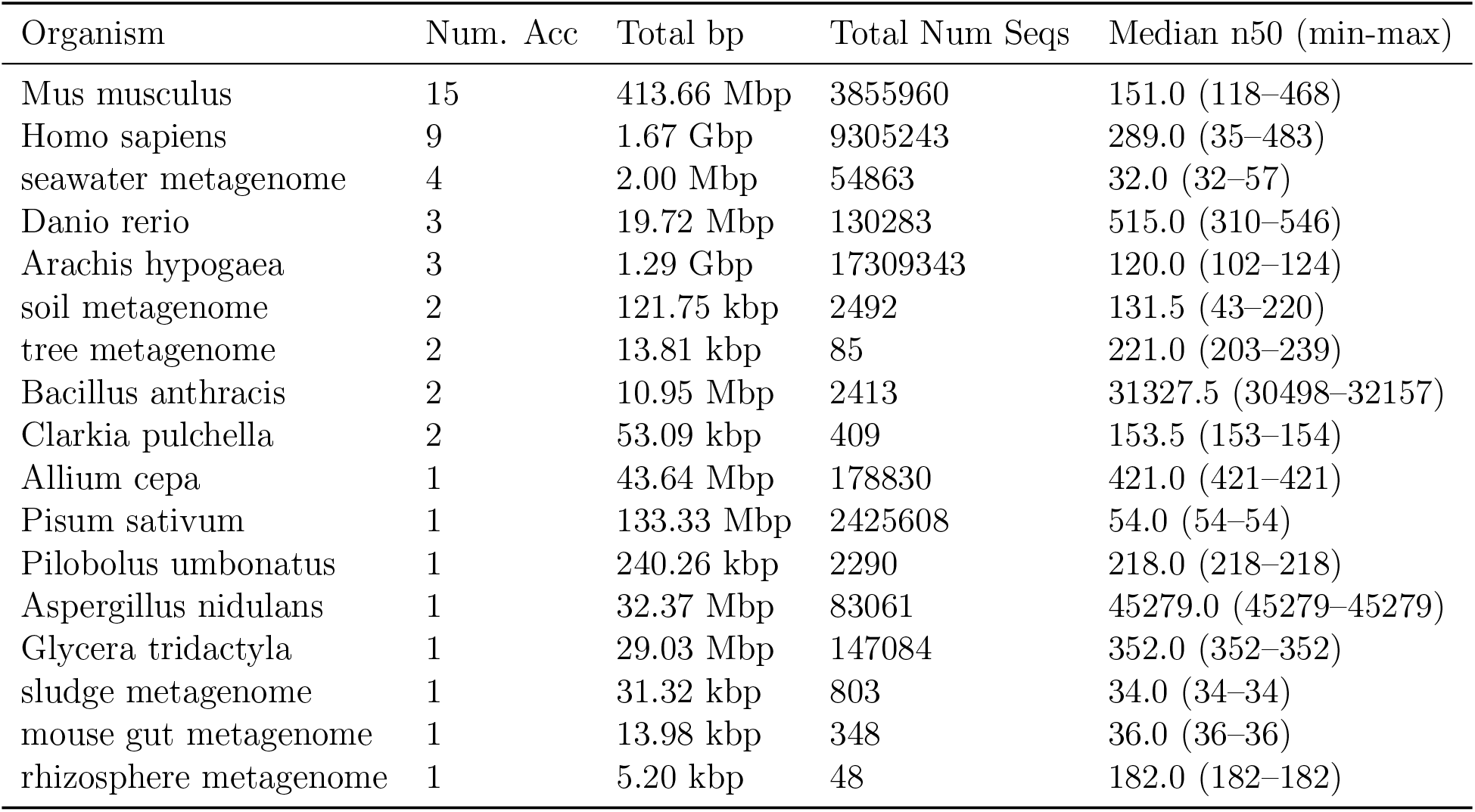
Training set composition of Logan training data.

| Organism | Num. Acc | Total bp | Total Num Seqs | Median n50 (min-max) |
| --- | --- | --- | --- | --- |
| Mus musculus | 15 | 413.66 Mbp | 3855960 | 151.0 (118–468) |
| Homo sapiens | 9 | 1.67 Gbp | 9305243 | 289.0 (35–483) |
| seawater metagenome | 4 | 2.00 Mbp | 54863 | 32.0 (32–57) |
| Danio rerio | 3 | 19.72 Mbp | 130283 | 515.0 (310–546) |
| Arachis hypogaea | 3 | 1.29 Gbp | 17309343 | 120.0 (102–124) |
| soil metagenome | 2 | 121.75 kbp | 2492 | 131.5 (43–220) |
| tree metagenome | 2 | 13.81 kbp | 85 | 221.0 (203–239) |
| Bacillus anthracis | 2 | 10.95 Mbp | 2413 | 31327.5 (30498–32157) |
| Clarkia pulchella | 2 | 53.09 kbp | 409 | 153.5 (153–154) |
| Allium cepa | 1 | 43.64 Mbp | 178830 | 421.0 (421–421) |
| Pisum sativum | 1 | 133.33 Mbp | 2425608 | 54.0 (54–54) |
| Pilobolus umbonatus | 1 | 240.26 kbp | 2290 | 218.0 (218–218) |
| Aspergillus nidulans | 1 | 32.37 Mbp | 83061 | 45279.0 (45279–45279) |
| Glycera tridactyla | 1 | 29.03 Mbp | 147084 | 352.0 (352–352) |
| sludge metagenome | 1 | 31.32 kbp | 803 | 34.0 (34–34) |
| mouse gut metagenome | 1 | 13.98 kbp | 348 | 36.0 (36–36) |
| rhizosphere metagenome | 1 | 5.20 kbp | 48 | 182.0 (182–182) |

**Table 9:**
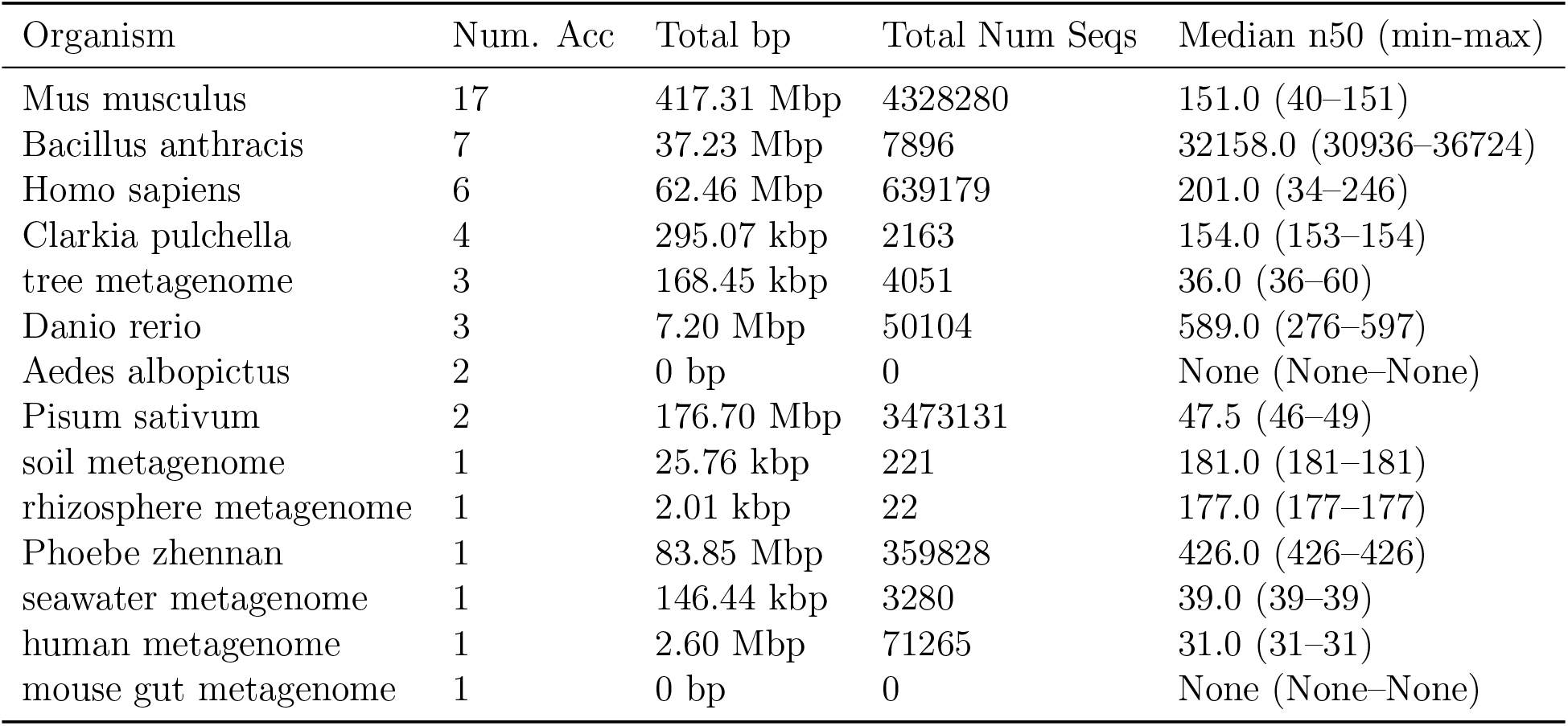
50 accession test set composition

**Table 10:**
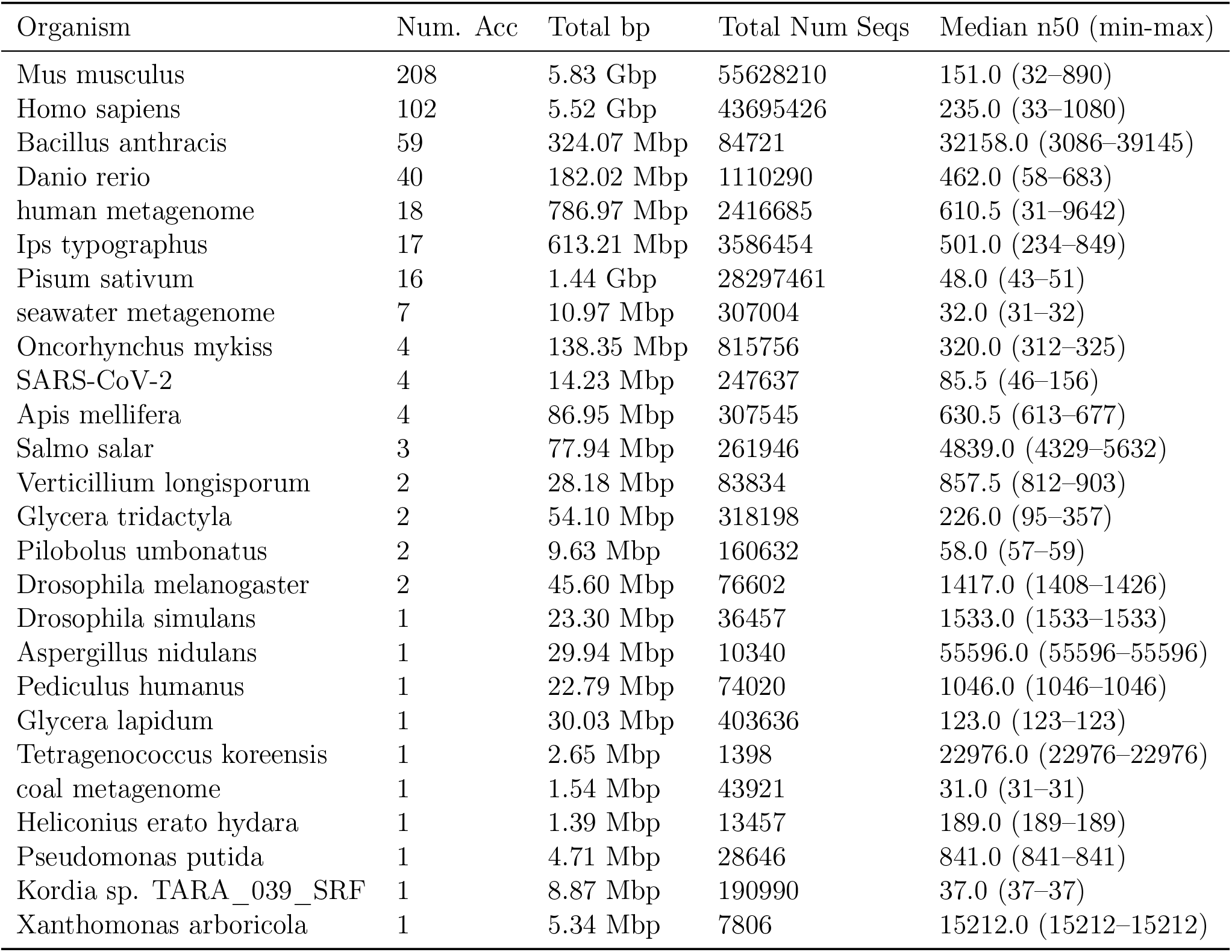
500 accession test set composition.

All accessions were selected as subsets from the roughly 5,000-accession 100-studies experiment from [21]. Training and testing accession sets were ensured to be exclusive.

To construct evaluation subsets, a target number of accessions are drawn by first sampling organisms with probability proportional to their frequency in the full 100-study pool of [21], then selecting one accession uniformly at random from that organism’s candidates. This procedure ensures that the organism composition of each subset mirrors that of the broader collection, capturing the taxonomic diversity of the original benchmark set.

#### A.2.3 Test Dataset Accessions

Information about the min and max base pair cutoffs for indexing and the sampling strategy

#### A.2.4 Reference Genome Training Dataset

Of the accessions in the Logan training data 8, there were 17 total "organisms". Of those, 6 were metagenomic accessions, leaving 11 species. 7 of those 11 species had genomes available for download on RefSeq, and they were downloaded to make up the reference sequence dataset. These collectively contained a total of 13 billion base pairs, which after chunking into 1024bp chunks left a total of 13.3M rows. These were randomly sampled down to 6 million to match the number of logan rows.

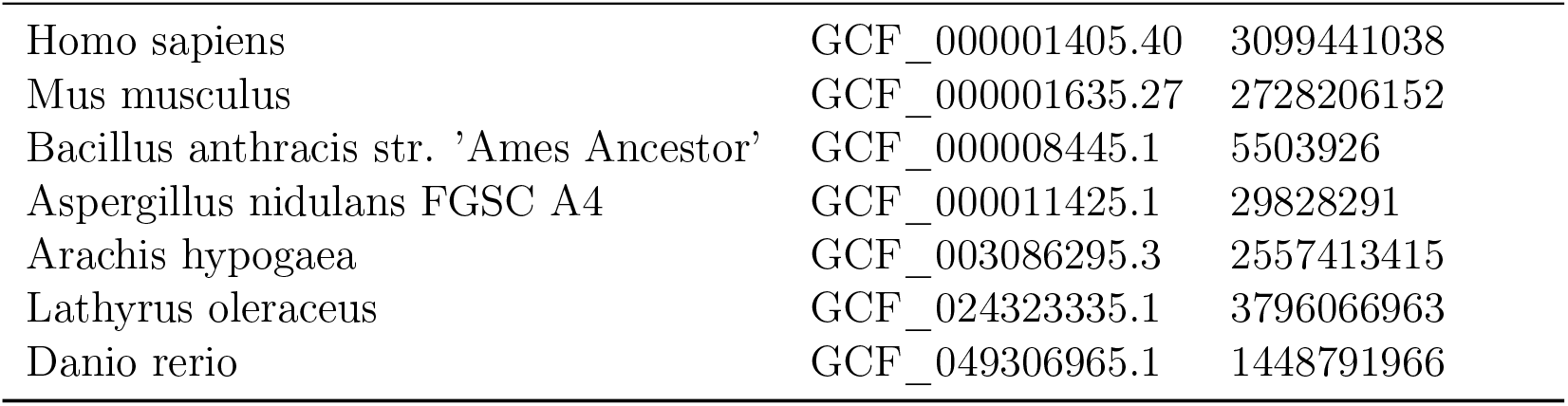

### A.3 Full Experimental Results

Tables 11, 12 and 13 give the full experimental results, with bootstrap standard errors, for the 50-, 500- and 4,571-accession experiments. IVF-PQ rows use the cuVS GPU index described in §4.4 with *m* = 100; the 1-bit row at 4,571 accessions is exhaustive RaBitQ search with *m* = 1000.

**Table 11:** 50-accession benchmark, all methods and variation rates. ± is the bootstrap standard error over 500 queries. Bold: best among scalable methods.

| Var. | Method | R@7 | R@$R_q$ | AUPRC |
| --- | --- | --- | --- | --- |
| 0% | LOCALE | 0.948±0.007 | 0.925±0.009 | 0.949±0.007 |
| | LOCALE + IVF-PQ | 0.874±0.012 | 0.823±0.014 | 0.861±0.012 |
| | MetaGraph | **0.984**±0.003 | **0.982**±0.005 | **0.983**±0.004 |
| | ESA | 0.954±0.007 | 0.936±0.008 | 0.953±0.007 |
| | LLM-ED | 0.886±0.011 | 0.785±0.015 | 0.836±0.012 |
| | MMseqs2 | 0.988±0.002 | 1.000±0.000 | 1.000±0.000 |
| 5% | LOCALE | **0.920**±0.009 | **0.850**±0.013 | **0.889**±0.011 |
| | LOCALE + IVF-PQ | 0.818±0.014 | 0.727±0.017 | 0.777±0.015 |
| | MetaGraph | 0.732±0.019 | 0.714±0.020 | 0.727±0.018 |
| | ESA | 0.792±0.014 | 0.610±0.018 | 0.688±0.015 |
| | LLM-ED | 0.757±0.016 | 0.552±0.019 | 0.640±0.016 |
| | MMseqs2 | 0.985±0.003 | 0.997±0.002 | 0.997±0.002 |
| 10% | LOCALE | **0.801**±0.015 | **0.632**±0.019 | **0.708**±0.015 |
| | LOCALE + IVF-PQ | 0.687±0.018 | 0.517±0.019 | 0.592±0.016 |
| | MetaGraph | 0.113±0.014 | 0.113±0.014 | 0.159±0.014 |
| | ESA | 0.510±0.019 | 0.251±0.016 | 0.360±0.014 |
| | LLM-ED | 0.617±0.018 | 0.343±0.018 | 0.448±0.015 |
| | MMseqs2 | 0.938±0.009 | 0.947±0.009 | 0.951±0.009 |

**Table 12:** 500-accession benchmark, all methods. ± is the bootstrap standard error over 500 queries. Bold: best among scalable methods. MMseqs2 (198 GB index, 80 s per 500-query batch on a 128-core CPU node) is the alignment reference.

| Var. | Method | R@10 | R@$R_q$ | AUPRC |
| --- | --- | --- | --- | --- |
| 0% | LOCALE | 0.882±0.011 | 0.876±0.012 | 0.901±0.010 |
| | LOCALE + IVF-PQ | 0.822±0.013 | 0.810±0.014 | 0.835±0.013 |
| | MetaGraph | **0.927**±0.009 | **0.954**±0.007 | **0.963**±0.006 |
| | ESA | 0.887±0.011 | 0.902±0.010 | 0.918±0.009 |
| | LLM-ED | 0.730±0.017 | 0.692±0.018 | 0.721±0.017 |
| | MMseqs2 | 0.935±0.008 | 0.993±0.003 | 0.994±0.003 |
| 5% | LOCALE | **0.808**±0.014 | **0.764**±0.015 | **0.796**±0.014 |
| | LOCALE + IVF-PQ | 0.727±0.016 | 0.675±0.017 | 0.704±0.016 |
| | MetaGraph | 0.686±0.020 | 0.654±0.020 | 0.659±0.019 |
| | ESA | 0.567±0.019 | 0.484±0.019 | 0.526±0.018 |
| | LLM-ED | 0.520±0.020 | 0.393±0.018 | 0.433±0.018 |
| | MMseqs2 | 0.934±0.008 | 0.990±0.004 | 0.993±0.003 |
| 10% | LOCALE | **0.569**±0.020 | **0.458**±0.019 | **0.507**±0.018 |
| | LOCALE + IVF-PQ | 0.480±0.019 | 0.400±0.019 | 0.437±0.018 |
| | MetaGraph | 0.099±0.013 | 0.097±0.013 | 0.106±0.013 |
| | ESA | 0.195±0.015 | 0.092±0.009 | 0.125±0.009 |
| | LLM-ED | 0.273±0.018 | 0.126±0.012 | 0.179±0.012 |
| | MMseqs2 | 0.895±0.011 | 0.941±0.010 | 0.944±0.009 |

**Table 13:**
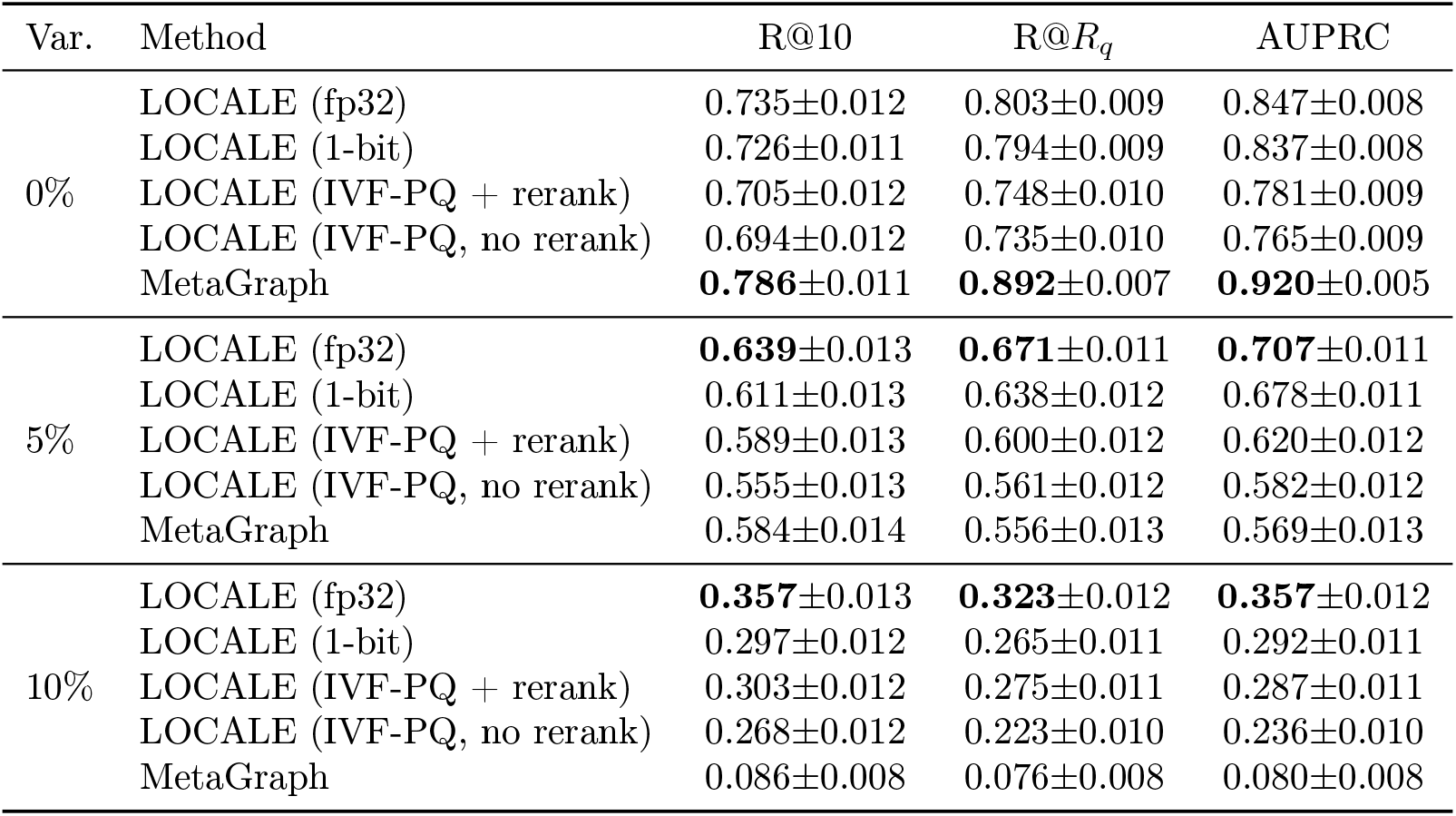
Full 4,571-accession benchmark (1,000 queries). Exact float32 and 1-bit RaBitQ search use *m* = 1000; IVF-PQ uses *m* = 100, probes 272 of 16,384 lists per shard, and optionally reranks the top 10 hits per shard with float32 vectors. ± is the bootstrap standard error. Bold: best per column.

| Var. | Method | R@10 | R@$R_q$ | AUPRC |
| --- | --- | --- | --- | --- |
| 0% | LOCALE (fp32) | 0.735±0.012 | 0.803±0.009 | 0.847±0.008 |
| | LOCALE (1-bit) | 0.726±0.011 | 0.794±0.009 | 0.837±0.008 |
| | LOCALE (IVF-PQ + rerank) | 0.705±0.012 | 0.748±0.010 | 0.781±0.009 |
| | LOCALE (IVF-PQ, no rerank) | 0.694±0.012 | 0.735±0.010 | 0.765±0.009 |
|  | MetaGraph | **0.786±0.011** | **0.892±0.007** | **0.920±0.005** |
| 5% | LOCALE (fp32) | **0.639±0.013** | **0.671±0.011** | **0.707±0.011** |
| | LOCALE (1-bit) | 0.611±0.013 | 0.638±0.012 | 0.678±0.011 |
| | LOCALE (IVF-PQ + rerank) | 0.589±0.013 | 0.600±0.012 | 0.620±0.012 |
| | LOCALE (IVF-PQ, no rerank) | 0.555±0.013 | 0.561±0.012 | 0.582±0.012 |
| | MetaGraph | 0.584±0.014 | 0.556±0.013 | 0.569±0.013 |
| 10% | LOCALE (fp32) | **0.357±0.013** | **0.323±0.012** | **0.357±0.012** |
| | LOCALE (1-bit) | 0.297±0.012 | 0.265±0.011 | 0.292±0.011 |
| | LOCALE (IVF-PQ + rerank) | 0.303±0.012 | 0.275±0.011 | 0.287±0.011 |
| | LOCALE (IVF-PQ, no rerank) | 0.268±0.012 | 0.223±0.010 | 0.236±0.010 |
| | MetaGraph | 0.086±0.008 | 0.076±0.008 | 0.080±0.008 |

#### A.3.1 Training Compute

##### Hyperparameters

We train with AdamW [26] (*β*_1_=0.9, *β*_2_=0.999, weight decay 10^−2^) at a peak learning rate of 6 × 10^−5^ with a 460-step linear warmup followed by cosine decay, with a per-device batch size of 64 across 4 nodes of 4 NVIDIA A100 GPUs (batch size 1024), for 5,859 steps, about 1 hour of wall-clock time (16 A100 hours). The approximate total compute used (tracked on a cluster account) for training totaled 961 node hours allocated, or 3844 A100 hours.

#### A.3.2 Data Cleaning Details of Section 4.3

This section details the methodology through which we chose the accessions tested in section 4.3.

Four genomes from genotypes A-D were chosen by querying the NCBI Nucleotide database for complete genomes of 3-3.3kb for each genotype-level organism of HBV using the [Organism] field. We selected the first hit of each, giving one genome per genotype as reference. See table 14 for a list of exact GenBank labels.

**Table 14:** Genotype reference genomes used for read-level screening. Selected from `NCBI nuccore` by querying `"hepatitis B virus genotype X"[Organism] AND complete genome[Title] AND 3000:3300[SLEN]`, taking one complete genome per genotype. The `[Organism]` constraint requires a genotype-level taxonomic assignment rather than submitter free text. Genotypes A and C are included so that the nearest-reference assignment can resolve to a genotype other than the one a run claims; only B and D appear in the benchmark arms.

| Genotype | Accession | Subgenotype | Length (bp) |
| --- | --- | --- | --- |
| A | LC851038.1 | A2 | 3,221 |
| B | LC851451.1 | B4 | 3,215 |
| C | LC886239.1 | C | 3,185 |
| D | LC852229.1 | D2 | 3,182 |

No genotype-D runs carry a genotype-level taxid on the SRA. Instead, to find reads we pulled all runs with the HBV organism `tax_tree(10407)` from the ENA (N=3850) and searched with regex `genotype[:\s]*([A-J])\b` over the following metadata fields:

- scientific_name
- strain
- isolate
- sample_title
- description

This left us with 423 total accessions (246 B, 81 D). We restricted to Illumina paired PCR runs. We chose Illumina to isolate biological divergence from sequencing error, PCR for whole-genome HBV at depth (rather than mostly host genome), and paired for consistency with the rest of the benchmark.

To ensure these raw reads actually corresponded to their labeled genotype, we performed a per-project label screening. We grouped runs by project, sampled 5-10 runs of each project and 2000 reads from each run, then mapped them to all references. For every read that mapped to the reference, we recorded its identity (number of matching bases over alignment block length) and took the median. For each sampled run, we assigned a label that was the reference with the maximum median identity score over all references. If any sampled run did not match its project’s label, we screened out the entire project.

For all projects that passed the label screen, we performed per-run screens. We screened read mapping rate, calculating the mapping rate to each reference genome and ensuring that the maximum over all references was ≥50%. We also screened breadth, ensuring that 75% of reference bases of the best matching reference were covered at least depth 10. We also filtered by those runs that were available on Logan, leaving 181 genotype B runs and 10 genotype D runs.

We took all 10 genotype D runs. We chose 5 genotype-B targets by sorting on coverage breadth and HBV read fraction and separating by project, choosing three from PRJNA556730 and two from PRJNA564199, all with breadth ≥ 0.995 and HBV fraction ≥0.80. See table 15 and 16 for the list of accessions that we used in our dataset.

**Table 15:** Genotype-B target accessions. Selected from the 181 QC-passing, Logan-indexed genotype-B runs on coverage breadth and HBV read fraction, with three drawn from PRJNA556730 and two from PRJNA564199 so that no single sequencing batch defines the relevant set. All candidates in this range have breadth ≥ 0.995 and HBV fraction ≥ 0.78, so the choice among near-ties is not material. Breadth is the fraction of reference positions covered at depth ≥ 10×.

| Accession | BioProject | Breadth | HBV frac. | Read len. |
| --- | --- | --- | --- | --- |
| SRR9843518 | PRJNA556730 | 1.0000 | 0.879 | 300 |
| SRR9843544 | PRJNA556730 | 0.9950 | 0.869 | 301 |
| SRR9843493 | PRJNA556730 | 0.9950 | 0.869 | 301 |
| SRR10076799 | PRJNA564199 | 0.9981 | 0.802 | 301 |
| SRR10076802 | PRJNA564199 | 0.9950 | 0.954 | 301 |

## B Varying The Returned Vector Shortlist Length

Search returns the top-*m* vectors per strand before accession-level aggregation (§3). Table 17 varies *m* on the 4,571-accession benchmark by regrouping the persisted exact float32 top-1000 shortlist at each depth. Recall@10 is flat in *m* because the best hit per accession is almost always within the first ten vectors, while Recall@*R_q_* and AUPRC, which depend on ranking accessions beyond the top ten, saturate between *m* = 100 and *m* = 200: *m* = 100 is within 0.5 points of *m* = 1000 at every variation rate, which motivates *m* = 100 for the approximate indexes.

**Table 16:** Genotype-D query accessions, all from PRJNA659786. All ten runs passing QC were used; no ranking or subselection was applied. Query and target accessions share no BioProject. Median read lengths are 131–149 bp against 300–301 bp in the target arm; queries are searched as read, without length equalization.

| Accession | Breadth | HBV frac. | Read len. |
| --- | --- | --- | --- |
| SRR12535931 | 0.9510 | 0.850 | 148 |
| SRR12535932 | 0.9620 | 0.857 | 149 |
| SRR12535933 | 0.9717 | 0.797 | 143 |
| SRR12535934 | 0.9733 | 0.829 | 140 |
| SRR12535938 | 0.8045 | 0.675 | 131 |
| SRR12535939 | 0.9110 | 0.779 | 146 |
| SRR12535940 | 0.8985 | 0.718 | 140 |
| SRR12535941 | 0.9268 | 0.799 | 131 |
| SRR12535945 | 0.9462 | 0.847 | 148 |
| SRR12535946 | 0.8164 | 0.721 | 132 |

**Table 17:** Shortlist depth *m* on the 4,571-accession benchmark (1,000 queries, exact float32 search, both strands). ± is the bootstrap standard error.

| $m$ | 0% | | | 5% | | | 10% | | |
| --- | --- | --- | --- | --- | --- | --- | --- | --- | --- |
| | $R@R_q$ | AUPRC | $R@10$ | $R@R_q$ | AUPRC | $R@10$ | $R@R_q$ | AUPRC | $R@10$ |
| 10 | 0.654 | 0.688 | 0.733 | 0.557 | 0.579 | 0.639 | 0.267 | 0.286 | 0.356 |
| 20 | 0.704 | 0.739 | 0.734 | 0.597 | 0.621 | 0.639 | 0.286 | 0.306 | 0.357 |
| 50 | 0.771 | 0.806 | 0.735 | 0.647 | 0.671 | 0.639 | 0.310 | 0.329 | 0.357 |
| 100 | 0.798 | 0.835 | 0.735 | 0.667 | 0.693 | 0.639 | 0.321 | 0.340 | 0.357 |
| 200 | 0.803 | 0.843 | 0.735 | 0.671 | 0.701 | 0.639 | 0.323 | 0.347 | 0.357 |
| 500 | 0.803 | 0.846 | 0.735 | 0.671 | 0.705 | 0.639 | 0.324 | 0.354 | 0.357 |
| 1000 | 0.803 | 0.847 | 0.735 | 0.671 | 0.707 | 0.639 | 0.323 | 0.357 | 0.357 |

Standard errors: 0.009–0.012 ( $R@R_q$ ), 0.008–0.012 (AUPRC), 0.011–0.013 ( $R@10$ ).

Table 18 asks how much an approximate index can miss before accession-level quality suffers. Starting from the same exact hits with *m* = 100, we randomly remove true top-100 vectors from each query’s shortlist so that the vector-level recall@100 equals a target value, and regroup. Vector recall of 0.99 costs under a point of AUPRC, 0.95 costs about 4 points, and 0.90 about 7 points; the deployed IVF-PQ index without rerank (0.765 AUPRC at 0%, Table 13) lands on the recall@100 ≈ 0.90 row, and the float32 rerank moves it to about 0.95.

**Table 18:** Accession-level quality as a function of the vector-level recall@100 of the shortlist, 4,571-accession benchmark, *m* = 100. The first row is exact search; the others subsample the exact top-100 hits to the stated vector recall. Standard errors 0.009–0.013.

| Vector recall@100 | 0% |  |  | 5% |  |  | 10% |  |  |
| --- | --- | --- | --- | --- | --- | --- | --- | --- | --- |
| | $R@R_q$ | AUPRC | $R@10$ | $R@R_q$ | AUPRC | $R@10$ | $R@R_q$ | AUPRC | $R@10$ |
| 1.00 (exact) | 0.798 | 0.835 | 0.735 | 0.667 | 0.693 | 0.639 | 0.321 | 0.340 | 0.357 |
| 0.99 | 0.792 | 0.829 | 0.730 | 0.662 | 0.687 | 0.634 | 0.320 | 0.337 | 0.355 |
| 0.95 | 0.760 | 0.793 | 0.699 | 0.636 | 0.656 | 0.606 | 0.305 | 0.323 | 0.344 |
| 0.90 | 0.735 | 0.765 | 0.680 | 0.615 | 0.632 | 0.585 | 0.296 | 0.311 | 0.333 |
| 0.80 | 0.677 | 0.702 | 0.633 | 0.564 | 0.576 | 0.542 | 0.273 | 0.281 | 0.303 |

## C Eval Dataset Stats

Table 19 summarizes the four evaluation bundles. Vectors are the number of embedded windows in the LOCALE index (encoder context 256 bp, 150 bp overlap). Queries are sampled raw reads after the relevance filters of §4; the 4,571-accession bundle holds 10,000 filtered queries, of which 1,000 are evaluated (seed 1337). |A*_q_*| is the number of relevant accessions per query.

## D Baseline Settings

### MMseqs2 settings

We run MMseqs2 [36] (commit d45e0c4) over a single sequence database containing every indexed accession’s Logan contigs, built with `createdb` followed by `createindex –search-type 3`. Each query batch is a single call to `easy-search` with `–search-type 3, –max-seqs 300` and `–format-output query,target,bits;` all other parameters are MMseqs2 defaults: sensitivity *s* = 5.7, seed length *k* = 15 with spaced *k*-mers, e-value cutoff 10^−3^, and no minimum sequence identity or coverage filter. The nucleotide search workflow extracts both query strands by default, so reverse-strand reads are found without any strand flag. Up to 300 prefilter candidates are aligned per query; every alignment passing the e-value cutoff is retained, and an accession is scored by the maximum bit score over its contigs. The reported search times are the mean of four repeats after a warm-up run, on one CPU node (two 64-core AMD EPYC 7763 processors) with `–threads 256`, and include query database creation, prefiltering, alignment and output conversion.

### D.1 Hardware

We measure search performance on a single node with one AMD EPYC 7763 64-core CPU, four NVIDIA A100 (Ampere) 40 GB GPUs, and 256 GB of DDR4 DRAM; the 4,571-accession IVF-PQ index is served from a node of the same design with four 80 GB A100s, since the index is resident in GPU memory. MetaGraph and MMseqs2 are CPU-only and run on a CPU node with two 64-core AMD EPYC 7763 processors and 512 GB of DRAM. Vector-index construction parallelizes across GPU nodes for embedding (4 nodes at 500 accessions, 32 at 4,571); exact float32 and 1-bit search at 4,571 accessions is sharded over 4 GPU nodes, while every other search, including IVF-PQ at 4,571 accessions, runs on a single node.

### D.2 Retrieval Quality

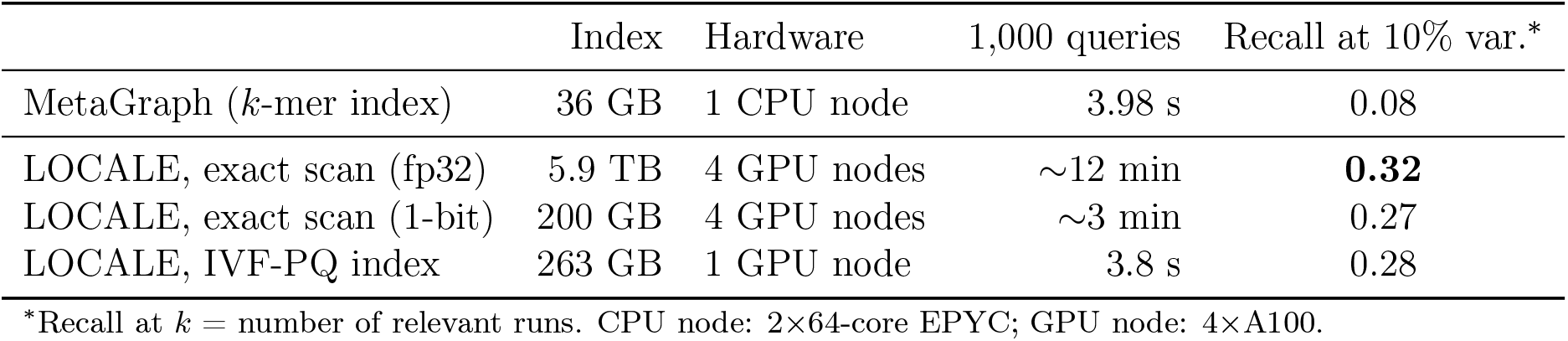

|  | Index | Hardware | 1,000 queries | Recall at 10% var.* |
| --- | --- | --- | --- | --- |
| MetaGraph ($k$-mer index) | 36 GB | 1 CPU node | 3.98 s | 0.08 |
| LOCALE, exact scan (fp32) | 5.9 TB | 4 GPU nodes | ~12 min | **0.32** |
| LOCALE, exact scan (1-bit) | 200 GB | 4 GPU nodes | ~3 min | 0.27 |
| LOCALE, IVF-PQ index | 263 GB | 1 GPU node | 3.8 s | 0.28 |

\*Recall at $k$ = number of relevant runs. CPU node: 2×64-core EPYC; GPU node: 4×A100.

**Table 19:** Evaluation dataset statistics. The HBV bundle is the 50-accession index plus the 5 genotype-B target accessions; its queries are genotype-D reads with a fixed relevant set of size 5, and it is evaluated without simulated variation.

|  | 50 acc. | HBV (55 acc.) | 500 acc. | 4,571 acc. |
| --- | --- | --- | --- | --- |
| Accessions indexed | 50 | 55 | 500 | 4,571 |
| Indexed contig sequence (Gbp) | 1.57 | 1.57 | 22.9 | 296 |
| Vectors in index (M) | 10.7 | 10.8 | 153.1 | 2,060 |
| Queries evaluated | 500 | 500 | 500 | 1,000 |
| Queries reverse-strand (%) | 55.2 | – | 51.0 | 53.4 |
| Mean query length (bp) | 128.3 | 118.5 | 135.9 | 137.5 |
| $\vert \mathcal{A}_q\vert $ mean / median / max | 2.51 / 1 / 10 | 5 / 5 / 5 | 5.21 / 2 / 59 | 17.2 / 3 / 385 |
| Simulated variation rates | 5%, 10% | none | 5%, 10% | 5%, 10% |
| Shortlist depth $m$ (exact) | 100 | 100 | 100 | 1,000 |

